# GENTANGLE: integrated computational design of gene entanglements

**DOI:** 10.1101/2023.11.09.565696

**Authors:** Jose Manuel Martí, Chloe Hsu, Charlotte Rochereau, Tomasz Blazejewski, Hunter Nisonoff, Sean P. Leonard, Christina S. Kang-Yun, Jennifer Chlebek, Dante P. Ricci, Dan Park, Harris Wang, Jennifer Listgarten, Yongqin Jiao, Jonathan E. Allen

## Abstract

**Summary:** The design of two overlapping genes in a microbial genome is an emerging technique for adding more reliable control mechanisms in engineered organisms for increased safety. The design of functional gene pairs is a challenging procedure and computational design tools are used to improve the efficiency to deploy successful designs in genetically engineered systems. GENTANGLE (Gene Tuples ArraNGed in overLapping Elements) is a high performance containerized pipeline for the computational design of two overlapping genes translated in different reading frames of the genome. This new software package can be used to design and test gene entanglements for microbial engineering projects using arbitrary sets of user specified gene pairs.

**Availability and Implementation:** The GENTANGLE source code and its submodules are freely available on GitHub at https://github.com/BiosecSFA/gentangle. The DATANGLE (DATA for genTANGLE) repository contains related data and results, and is freely available on GitHub at https://github.com/BiosecSFA/datangle. The GENTANGLE repository wiki contains detailed instructions on how to use the container and the different components of software and data, including reproducing the results. The code is licensed under the GNU Affero General Public License version 3 (https://www.gnu.org/licenses/agpl.html).

## 1 Introduction

We present GENTANGLE, a pipeline for enhanced computational design of overlapping genes to prolong synthetic device function and limit horizontal gene transfer. When engineered bacteria with synthetic biology designs are used in natural environments, their engineered functions can be lost rapidly due to genetic instability or can be horizontally acquired by native bacteria (Kumar and Hasty, 2023). One strategy to prolong the desired function and limit horizontal gene transfer (HGT) is to design two genes in overlapping reading frames, referred to as “gene entanglement”, which can be partial or complete (Arbel-Groissman *et al*., 2023). An example of the partial approach is when the two reading frames overlap, but not the actual coding sequences of the corresponding pair of proteins (Decrulle *et al*., 2021). Recently published software, CAMEOS (Constraining Adaptive Mutations using Engineered Overlapping Sequences) (Blazejewski *et al*., 2019), written in Julia (Bezanson *et al*., 2017), was introduced to design fully overlapping genes. However, CAMEOS is a computationally intensive sequential code requiring several pre-computed inputs. More recently, an effort was made to distribute CAMEOS in a virtual machine to expand accessibility to the original software (Logel and Jaschke, 2023). Nevertheless, the original code remains limited in the ability to efficiently evaluate new gene pairs (Arbel-Groissman *et al*., 2023). To automate the entire process of gene entanglement, enhance its computational scalability, improve software portability, and add new functional and analysis capabilities, we developed GENTANGLE, a computational biology pipeline built around CAMEOX (CAMEOs eXtended), an enhanced, parallelized version of CAMEOS that we release as part of GENTANGLE.

## 2 Features

GENTANGLE provides an end-to-end computational pipeline for the design of gene entanglements. In GENTANGLE, the entire sequence entanglement design process has been automated and integrated into a high performance pipeline, deployed using a single Singularity container to support portability and reproducible output. This container exposes more than ten entry points to the user, adhering to the Scientific Filesystem (SCIF) standard (Sochat, 2018) for encapsulating multiple applications into one container. The use of a single container based on a SCIF-compliant recipe for a complex pipeline significantly increases the robustness and maintainability of the software.

The GENTANGLE pipeline (Figure 1A) is composed of three main sections, 1) upstream: protein sequence preparation for fitness modeling, core: protein fitness estimation and sequence entanglement search (CAMEOX), and 3) downstream: analysis and visualization of gene entanglement solutions. CAMEOX adds multi-thread parallelism and a dynamic stopping criteria to the previously published CAMEOS software, enabling substantial speed-ups. When run in a multi-threaded environment, speed-ups can increase beyond an order of magnitude. In CAMEOX, each entanglement candidate sequence is modified and scored for predicted fitness over a different number of iterations. Unlike the original CAMEOS software, the change in fitness score is tracked and an automated stopping criteria is invoked to terminate the search when the fraction of candidates that sustain a change in fitness score decreases below a user defined threshold. This allows the software to increase the number of coordinated runs in parallel by efficiently skipping candidate sequence iterations that have converged to their locally optimal fitness score. The upstream section of the pipeline generates all fitness modeling inputs for each protein pair from multiple sequence alignments (MSAs) curated via EVcouplings (Hopf *et al*., 2018), including training hidden Markov models (HMM) with HMMER (Mistry *et al*., 2013) and Markov random field (MRF) models with CCMpred (Ekeberg *et al*., 2013). Since the protein fitness models are derived from evolutionary sequence conservation patterns of an individual protein family, high-quality MSAs are needed. To accommodate diverse gene family types, the GENTANGLE pipeline offers two different methods for building MSAs. One method is based on UniProt’s UniRef90 or UniRef100 (Consortium, 2022) to allow for a larger set of input sequences, which is advantageous for training MRF models (Blazejewski *et al*., 2019). While this has the benefit of building a fitness model derived from a larger set of example sequences, it may inadvertently include conserved sequences with a divergent function. An alternative method uses OrthoDB to start with a curated set of gene orthologs to restrict inclusion of genes with a functional match to the target gene (Kriventseva *et al*., 2019), while accepting the potential for fewer training examples.

**Fig. 1.**
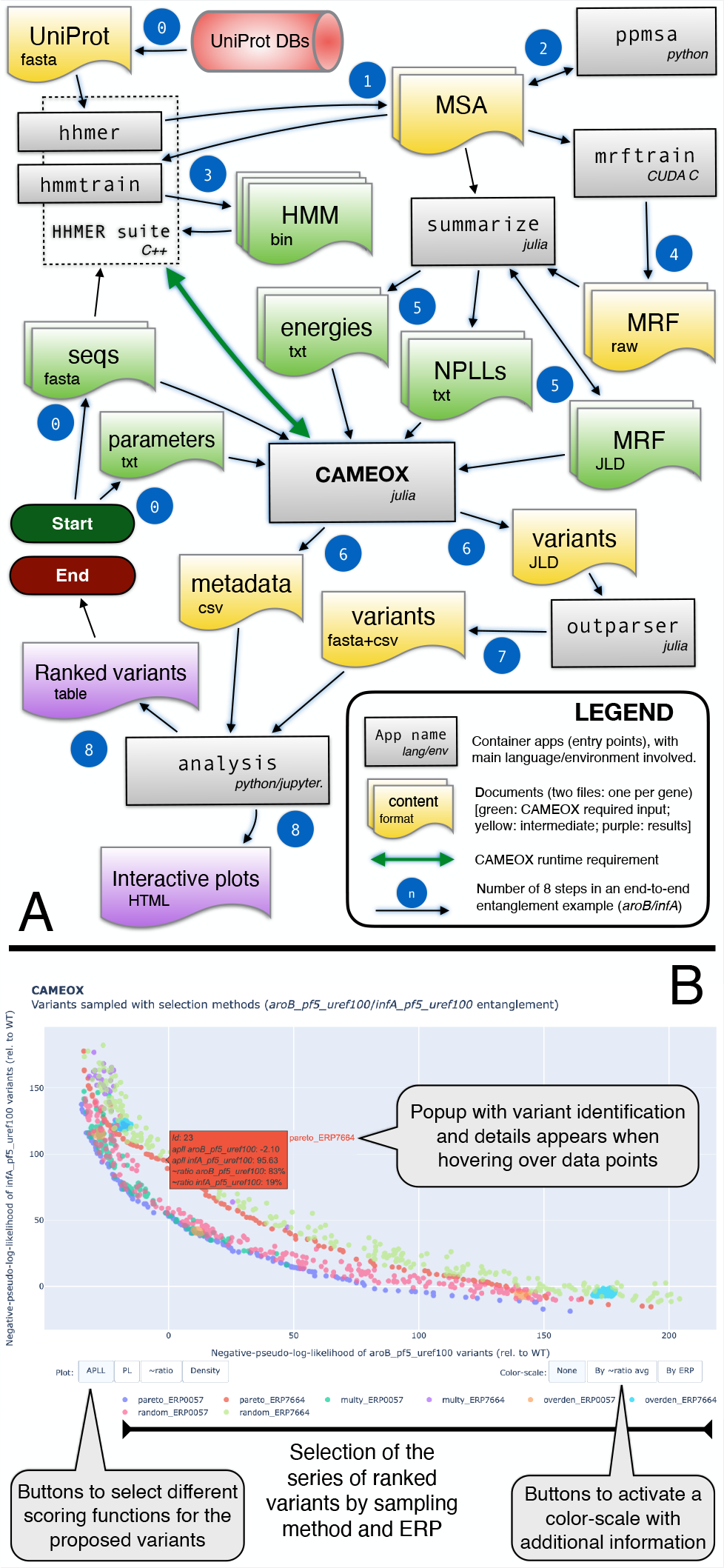
(A) GENTANGLE is a containerized computational pipeline for sequence entanglement. The upstream pipeline generates all the inputs required by CAMEOX, the parallel code proposing entangled variants. This includes training HMM and MRF models for each protein pair after obtaining curated multi-sequence alignments. The downstream pipeline produces quantitative analyses and interactive visualization of CAMEOX results to inform the selection of candidate variants. This flowchart depicts the main steps for an end-to-end gene entanglement example (*aroB* and *infA* on *Pseudomonas* PF5 as host) as detailed in the GENTANGLE wiki at https://github.com/BiosecSFA/gentangle/wiki. Each of the numbered steps corresponds to different apps within the singularity image, which are distinct points of entry to the container that follow the SCIF standard. Step 0 is a preparatory step (data download) that is detailed in the wiki. This example uses the data in the DATANGLE repository of example data for GENTANGLE, which is freely available on GitHub at https://github.com/BiosecSFA/datangle. **(B) Results of selected variants presented in an interactive plot**. This example snapshot presents one of the three interactive plots that the “analysis” container app generates as part of the final results, in addition to different files in tabular format. The figure highlights the different controls for interacting with the variants (generic navigation controls not shown), such as the buttons to activate a color-scale to include the averaged similarity with the WT sequences or the ERP. The scatter plot shows the selected variants for the four sampling criteria (pareto, multiplicity, overdensity, and random) for two different ERP (0.0057 and 0.7664).

New functional capability is included in CAMEOX to efficiently explore gene entanglements in new organisms and genes of interest. Users can now specify codon usage tables (Athey *et al*., 2017) obtained via CoCoPUTS (Alexaki *et al*., 2019) for design projects in organisms outside of *Escherichia coli* by indicating its NCBI taxonomic identifier (Federhen, 2011). The software allows the user to adjust the influence of the fitness score of one gene over its partner. Instead of assuming an equal contribution for the fitness score of each gene pair, one gene can be weighted more heavily or a random weight can be assigned to each gene. Adjusting the weight of each gene and its partner changes the combined entanglement score and can influence the explored space of candidate solutions. Lastly, the user can now increase the number and length of mutations introduced into candidate entanglements to more effectively explore a larger variation of the candidate sequence space. In the original CAMEOS, each seed entanglement sequence is re-scored by the MRF fitness models by iteratively introducing single mutations as the score improves. Allowing multiple mutations to be introduced in a single iteration gives the option to move to more divergent parts of the sequence space at each iteration and possibly expand the sequence diversity of solutions.

The GENTANGLE pipeline includes newly developed software to visualize and select CAMEOX sequence proposals. The new pyCAMEOX python module is used as a back-end to a Jupyter notebook front-end. In addition to running the notebook interactively, the user may choose to perform a parameterized automated analysis using a dedicated point of entry in the GENTANGLE singularity container. The solution space is analyzed using pandas (Wes McKinney, 2010; Pandas dev team, 2022) and visualized with interactive HTML files automatically generated using matplotlib (Hunter, 2007) and plotly (P.T.Inc., 2015) graphics libraries (see Figure 1B for an example of interactive scatter plot). Each candidate solution plots the negative pseudo loglikelihood (NPLL) scores predicting the fitness potential of each protein in the entangled gene. Additional information for each solution includes sequence similarity between the synthetic sequence and wild type, and the relative starting position of the shorter gene embedded in the longer gene referred to as the Entanglement Relative Position (ERP). Solutions can be ranked and selected according to three criteria in addition to random sampling of proposed variants: 1) an ordered list of solutions along the Pareto frontier, 2) ranking by multiplicity of the solution —different HMM seeds converging to the same solution after the MRF optimization, and 3) ranked sampling from different overdensities appearing in the NPLL solution space. Each criterion reports a ranked list of variants with the selected proposals ordered by their score using the particular selection strategy. The sampled variants are saved in different files and presented in an interactive plot (Figure 1B). The criteria can be combined in a multi-criterion sampling-without-replacement strategy. For instance, the multiplicity criterion returns as the top variant the most repeated solution among all the CAMEOX runs considered for a given entanglement pair. For the overdensity criterion, the NPLL space is searched for a tentative number of non-overlapping ranges corresponding to a higher density of variants while maximizing the pairwise distance of the range’s centers of mass. The NPLL scores are initially grouped into discrete bins with similarly scored solutions with the goal of making a balanced selection of proposed solutions across the span of predicted fitness values. To pick variants from the selected ranges, a round-robin ranked sampling is used to choose a balanced number of variants that are ranked based on the approximate local density. The developed algorithms rely on different packages of SciPy (Virtanen *et al*., 2020) and NumPy (Harris *et al*., 2020). This general approach maybe valuable at early design phases where confidence in the fitness predictions is not well understood. The different selection criteria are made available to allow users to explore different parts of the solution space depending on confidence in the underlying fitness models. In addition, the synthetic sequence proposals can be clustered and visualized with external tree clustering software, with output tested for ETE3 (Huerta-Cepas *et al*., 2016) and PhyloCloud (Deng *et al*., 2022).

## 3 Conclusion

The complexity of designing entangled gene pairs presents a practical barrier for its use as part of a control strategy for engineered organisms. Although, recent work has shown the potential to produce functional entanglements in an environmentally relevant microbe (Chlebek *et al*., 2023), current progress is impaired by the lack of flexible and efficient computational tools to allow microbial engineers to effectively test new designs. GENTANGLE makes available for the first time an efficient software tool that can be applied to arbitrary gene pairs to expand the opportunity to computationally propose and experimentally test new entanglement designs. Developing reliable control mechanisms for genetically modified organisms will be critical to a successful expansion of synthetic biology.

## References

Alexaki, A. et al. (2019). Codon and codon-pair usage tables (cocoputs): Facilitating genetic variation analyses and recombinant gene design. Journal of Molecular Biology, 431(13), 2434–2441. Computation Resources for Molecular Biology.

Arbel-Groissman, M. et al. (2023). Fighting the battle against evolution: designing genetically modified organisms for evolutionary stability. Trends in Biotechnology, 41, 1518–1531.

Athey, J. et al. (2017). A new and updated resource for codon usage tables. BMC bioinformatics, 18(1), 1–10.

Bezanson, J. et al. (2017). Julia: A fresh approach to numerical computing. SIAM review, 59(1), 65–98.

Blazejewski, T. et al. (2019). Synthetic sequence entanglement augments stability and containment of genetic information in cells. Science, 365(6453), 595–598.

Chlebek, J. L. et al. (2023). Prolonging genetic circuit stability through adaptive evolution of overlapping genes. Nucleic Acids Research, 51(13), 7094–7108.

Consortium, T. U. (2022). UniProt: the Universal Protein Knowledgebase in 2023. Nucleic Acids Research. gkac1052.

Decrulle, A. L. et al. (2021). Engineering gene overlaps to sustain genetic constructs in vivo. PLOS Computational Biology, 17(10), 1–19.

Deng, Z. et al. (2022). PhyloCloud: an online platform for making sense of phylogenomic data. Nucleic Acids Research, 50(W1), W577–W582.

Ekeberg, M. et al. (2013). Improved contact prediction in proteins: Using pseudolikelihoods to infer potts models. Phys. Rev. E, 87, 012707.

Federhen, S. (2011). The NCBI Taxonomy database. Nucleic Acids Research, 40(D1), D136–D143.

Harris, C. R. et al. (2020). Array programming with NumPy. Nature, 585(7825), 357–362.

Hopf, T. A. et al. (2018). The EVcouplings Python framework for coevolutionary sequence analysis. Bioinformatics, 35(9), 1582–1584.

Huerta-Cepas, J. et al. (2016). ETE 3: Reconstruction, Analysis, and Visualization of Phylogenomic Data. Molecular Biology and Evolution, 33(6), 1635–1638.

Hunter, J. D. (2007). Matplotlib: A 2d graphics environment. Computing in Science & Engineering, 9(3), 90–95.

Kriventseva, E. V. et al. (2019). Orthodb v10: sampling the diversity of animal, plant, fungal, protist, bacterial and viral genomes for evolutionary and functional annotations of orthologs. Nucleic acids research, 47(D1), D807–D811.

Kumar, S. and Hasty, J. (2023). Stability, robustness, and containment: preparing synthetic biology for real-world deployment. Current Opinion in Biotechnology, 79, 102880.

Logel, D. Y. and Jaschke, P. R. (2023). Creating De Novo Overlapped Genes, pages 95–120. Springer US, New York, NY.

Mistry, J. et al. (2013). Challenges in homology search: HMMER3 and convergent evolution of coiled-coil regions. Nucleic Acids Research, 41(12), e121–e121.

Pandas dev team (2022). pandas-dev/pandas: Pandas.

P.T.Inc. (2015). Collaborative data science.

Sochat, V. (2018). The Scientific Filesystem. GigaScience, 7(5). giy023.

Virtanen, P. et al. (2020). SciPy 1.0: Fundamental Algorithms for Scientific Computing in Python. Nature Methods, 17, 261–272.

Wes McKinney (2010). Data Structures for Statistical Computing in Python. In Stéfan van der Walt and Jarrod Millman, editors, Proceedings of the 9th Python in Science Conference, pages 56–61.

